# Aspirin hastens resolution of skeletal muscle inflammation and promotes recovery of muscle strength following acute injury

**DOI:** 10.64898/2026.04.21.719989

**Authors:** Xinyue Lu, Hamood Rehman, Amelie S. Sercu, James F. Markworth

## Abstract

Nonsteroidal anti-inflammatory drugs (NSAIDs) are widely recognized to potentially interfere with skeletal muscle regeneration. However, current knowledge is based almost exclusively on non-aspirin NSAIDs. Aspirin (ASA) differs from other NSAIDs in its ability to irreversibly acetylate cyclooxygenase-2 (COX-2), thereby redirecting its activity toward a lipoxygenase (LOX)-like function that enables the production of unique ASA-triggered specialized pro-resolving lipid mediators (AT-SPMs). Despite this, the potential impact of ASA on musculoskeletal tissue repair remains poorly understood. This study directly compared the effect of ASA against non-ASA NSAIDs on *in vitro* myogenesis and *in vivo* skeletal muscle injury and regeneration. Unlike non-ASA NSAIDs, including indomethacin (INDO), celecoxib, and SC-236, which markedly impaired C2C12 myotube formation at concentrations near their pharmacological ranges, ASA only interfered with myogenesis at overtly supraphysiological concentrations. In mice, an oral dose of 3 mg/kg/day INDO following barium chloride-induced muscle injury reduced regenerating myofiber cross-sectional area and impaired the recovery of muscle force-generating capacity. In contrast, a potency-matched oral treatment with 30 mg/kg/day ASA hastened the resolution of cellular inflammation, promoted myonuclear accretion, and improved recovery of absolute muscle strength. The beneficial effects of ASA on inflammatory resolution and muscle strength—but notably not myonuclear accretion—were reversed in mice co-treated with ASA + INDO. These findings demonstrate that, unlike non-ASA NSAIDs, ASA does not impair skeletal muscle regeneration and may promote a favorable early inflammatory environment for repair via unique COX-dependent pro-resolving and COX-independent anabolic mechanisms.

## INTRODUCTION

Non-steroidal anti-inflammatory drugs (NSAIDs) are commonly used to alleviate pain, swelling, and redness following acute and chronic musculoskeletal tissue injury^1^. NSAIDs exert their anti-inflammatory and analgesic effects by inhibiting the cyclooxygenase-1 and −2 (COX-1 and −2) enzymes, which catalyze the conversion of the long-chain (LC) omega-6 (n-6) polyunsaturated fatty acid (PUFA) arachidonic acid (ARA, 20:4n-6) to form prostaglandin H_2_ (PGH_2_), the precursor for major prostaglandins including PGD_2_, PGE_2_, PGF_2α_, PGI_2_ (prostacyclin), and TXA_2_ (thromboxane)^2^.

Prostaglandins are produced locally in response to tissue damage and function as autocrine/paracrine signaling molecules to promote vasodilation, stimulate leukocyte chemotaxis, and sensitize local nociceptors^3^. However, numerous studies have provided evidence to suggest that NSAID treatment may inadvertently interfere with the ability of damaged skeletal muscle to regenerate successfully^4^. This may be due to both deregulation of immune-muscle cell crosstalk^5^, as well as direct suppressive effects of NSAIDs on myogenesis^6^. Indeed, specific prostaglandins have been shown to be directly involved in various steps and the myogenic program. For example, PGE_2_ is critical for the proliferation of muscle stem cells^7–10^, PGI_2_ promotes metabolic reprogramming via its IP receptor^11–13^, PGD_2_ enhances myoblast proliferation and modulates fusion timing^14–16^, and PGF_2α_ facilitates muscle cell survival^17^, fusion^18^ and hypertrophy^19^.

Aspirin (acetylsalicylic acid; ASA) is one of the oldest and most consumed drugs in the world, with more than 50 million regular users in the United States alone^20^. While originally identified as a founding member of the NSAID class^21^, ASA is unique in that unlike other NSAIDs it acts to irreversibly acetylate the COX-1 and −2 isoforms^22^. Acetylation of Ser529 in COX-1 by ASA renders the enzyme completely inactive, thereby preventing prostaglandin biosynthesis^23^. Interestingly, however, acetylation of Ser516 in COX-2 by ASA does not completely inhibit its enzymatic activity but rather results in a switch from COX activity to a 15-lipoxygenase (15-LOX)-like function^24,25^.

Endogenous mammalian LOX enzymes, including 5-, 12-, and 15-LOX, function to oxygenate the 20-carbon ARA substrate at specific sites to form 5-, 12-, and 15-hydroxy-eicosatetraenoic acids (HETEs)^26^. Similarly, when using the 22-carbon omega-3 (n-3) PUFA docosahexaenoic acid (DHA, 22:6, n-3) as a substrate, these LOX enzymes produce 7-, 14-, and 17-hydroxy-docosahexaenoic acids (HDoHEs), respectively^27^. Such monohydroxylated PUFA metabolites formed via the standard LOX pathways carry their hydroxy groups in the S-configuration. In contrast, ASA-acetylated COX-2 functions to produce 15(R)-HETE^28^ and 17(R)-HDoHE^29^ enantiomers from ARA and DHA substrates, respectively. Endogenously produced 15(S)-HETE and 17(S)-HDoHE are important precursors in the downstream biosynthesis of di- and tri-hydroxylated specialized pro-resolving mediators (SPMs) via the 5-LOX pathway, including the lipoxins (e.g., LXA_4_) and D-series resolvins (e.g., RvD1)^30^. Similarly, 15(R)-HETE and 17(R)-HDoHE produced by ASA-acetylated COX-2 can be converted via the action of 5-LOX to form unique ASA-triggered (AT) lipoxins (e.g., AT-LXA4)^31^, resolvins (e.g., AT-RvD1 and AT-RvD3)^29^, and neuroprotectins (e.g., NPD1)^32^. While specific R-epimeric ‘AT’ labels are exclusive to the lipoxins and D-series resolvins, the entire E-series resolvin family (e.g., RvE1 and RvE2) derived from n-3 eicosapentaenoic acid (EPA, 20:5n-3) is also classically recognized as being initiated by this ASA-triggered switch^33^. Collectively, this body of literature suggests that unlike non-ASA NSAIDs, which can be resolution-toxic via their suppressive effects on endogenous resolution circuits^34,35^, ASA may uniquely expedite the resolution of inflammation by increasing AT-SPM production^36^.

Most studies on skeletal muscle growth and regeneration have focused on non-ASA NSAIDs, including common non-selective agents like ibuprofen (IBU)^37–43^ and indomethacin (INDO)^7,44–48^, as well as selective COX-2 inhibitors like NS-398^49–52^, SC-236^53–55^, and celecoxib^56–58^. Although a limited number of studies have tested the effect of ASA on skeletal muscle, it has typically been used as a representative non-selective COX inhibitor^59–63^, or as a presumed inactive placebo control^64,65^, rather than the main variable of interest. Consequently, the potential unique mechanisms of action of ASA, such as the production of AT-SPMs via the COX-2 acetylation switch, remain largely unexplored in the physiological context of muscle injury and repair.

Recent studies by us and other groups have shown that pharmacological administration of synthetic forms of endogenous S-enantiomer forms of SPMs, including RvD1^66,67^, RvD2^68–70^, PD1^71^, and MaR1^72^ can expedite resolution of the inflammatory response and stimulate tissue regeneration following acute skeletal muscle injury. Consistently, transgenic (*Alox15*^-/-^) mice that lack the endogenous murine 12/15-LOX enzyme, the primary biosynthetic enzyme for generating endogenous S-enantiomer precursors for lipoxins, resolvins, and maresins, mount an exaggerated acute inflammatory response to skeletal muscle injury in association with an intramuscular imbalance of pro-inflammatory vs. pro-resolving lipid mediators^73^. These mice also display impaired myofiber regenerative capacity, including defects in myogenic gene expression and myofiber size^71,73^. Indeed, pharmacological or genetic ablation of 12/15-LOX was recently discovered to markedly block *in vitro* myogenesis even in the absence of a functional host immune system demonstrating that *Alox15* plays a novel direct and indispensable role in determination of myogenic cell fate^71,73^.

Expanding beyond endogenous SPM circuits, one recent study showed that local hydrogel-based delivery of the 17R-ASA-triggered RvD1 epimer (AT-RvD1) enhanced muscle regeneration, improved muscle function, and reduced pain following volumetric muscle loss in mice^74^. However, to our knowledge no study to date has tested the potential ability of ASA as a cheap and readily available tool to potentially increase endogenous biosynthesis of AT-SPMs and the resulting downstream impact on inflammation and regeneration following muscle injury.

The purpose of the present study was to determine the effects of ASA on the temporal coordination of skeletal muscle regeneration following acute injury. Utilizing a barium chloride (BaCl_2_)-induced model of myofiber necrosis in mice, we conducted a side-by-side comparison between ASA and INDO to distinguish between ASA-triggered resolution pathways and potent, non-selective COX inhibition. We hypothesized that while INDO would display resolution-toxic effects by blunting the necessary inflammatory-to-repair transition, ASA would hasten resolution of acute inflammation and promote superior myofiber regeneration. Since AT-SPM biosynthesis requires acetylated COX-2 activity, we further hypothesized that the beneficial effects of ASA would be reversed in mice receiving combinational treatment with INDO + ASA.

## MATERIALS AND METHODS

### Muscle Cell Culture

The mouse C2C12 skeletal muscle cell line (ATCC, CRL-1772) was cultured in high glucose Dulbecco’s modified Eagle medium (DMEM, Gibco, 11995-065) supplemented with 10% fetal bovine serum (FBS) (Corning, 35-010-CV) and antibiotics (penicillin 100 U/mL, streptomycin 100 μg/mL) (Gibco, 15140-122) at 37 °C in a 5% CO_2_ atmosphere. C2C12 myoblasts were seeded into 12-well culture plates at a cell density of 2.5 × 10^4^/cm^2^ and allowed to proliferate and crowd in growth media for 72 hours. To induce myogenic differentiation, the growth media of confluent C2C12 myoblasts was replaced by differentiation media consisting of DMEM supplemented with 2% horse serum (HS, Gibco 26050088) and antibiotics. At the onset of induction of myogenic differentiation, cells were treated with ASA (Cayman Chemical, 70260, 62.5 µM**–**4 mM), common non-selective NSAIDs including INDO (Cayman Chemical, 70270, 6.25**–**400 µM) and IBU (Cayman Chemical, 70280, 1 µM), or COX-2 selective NSAIDs (COXIBs) including celecoxib (Cayman Chemical, 10008672, 50 µM) and SC-236 (Cayman Chemical, 10004219, 25 µM). NSAID stock solutions were prepared in 100% ethanol and stored at −20°C. Control cultures were treated differentiation media containing with the matching ethanol vehicle (≤0.8%). Differentiating C2C12 myoblasts were cultured in the continued presence of respective NSAIDs without a media change for 72 hours before fixation and immunocytochemistry as described below.

### Immunocytochemistry

C2C12 myotubes were fixed in 4% paraformaldehyde (PFA, Electron Microscopy Sciences, 15710) for 30 minutes at 4°C and then permeabilized with 0.1% Triton X-100 for 30 minutes at room temperature. The cells were then blocked in 1% bovine serum albumin (BSA, Sigma-Aldrich A3294-50G) for 1 hour at room temperature prior to overnight incubation at 4°C with primary antibodies against sarcomeric myosin (MF20c, DSHB, 1:100) and myogenin (F5Dc, DSHB, 1:100) prepared in blocking buffer. The following morning, cells were washed 3 × 5 min in PBS and then incubated for 1 hour at room temperature with PBS containing a mixture of Alexa Fluor conjugated secondary antibodies including Goat Anti-Mouse IgG2b Alexa Fluor 647 (Invitrogen, Thermo Fisher Scientific A-21242, 1:500) and Goat Anti-Mouse IgG1 Alexa Fluor 568 (Invitrogen A-21124, Thermo Fisher Scientific, 1:500), together with DAPI (Invitrogen, Thermo Fisher Scientific D21490, 2 μg/mL) to counterstain cell nuclei. Cells were washed with PBS at least three times before imaging. Stained C2C12 cells were visualized using an automated fluorescent microscope operating in inverted configuration (Echo Revolution). Nine images per well were automatically captured from the same predetermined locations using a 10 × Plan Fluorite objective. Total cell number (DAPI^+^ cells/mm^2^), myosin-positive (myotube) area (μm^2^) per field of view, and differentiation index (% DAPI nuclei within myosin^+^ cytoplasm) were quantified using a custom automated in-house plugin for ImageJ/FIJI.

### Mouse Skeletal Muscle Injury Model

Female C57BL/6NCrl mice (6-8 months old) were purchased from Charles River Laboratories and housed under specific pathogen-free (SPF) conditions with ad libitum access to food and water. Mice were anesthetized with 2% isoflurane and received unilateral intramuscular injection of the left TA muscle with 50 μL of 1.2% barium chloride (BaCl_2_) dissolved in sterile saline. Mice were returned to their home cage to recover and monitored until ambulatory. To assess the extent of inflammation and regeneration, TA muscles were collected on day 1, day 3, day 5, and day 14 following muscle injury. A minimum of n=5 mice (biological replicates) were included in each experimental group.

### Oral Administration of NSAIDs to Mice

United States Pharmacopeia (USP) grade INDO and ASA in powder form were obtained from the Purdue University Veterinary Pharmacy. INDO and ASA were dissolved in 100% ethanol at concentration of 7.5 and 75 mg/mL, respectively and stored short term at −20°C. On the day of treatment, INDO and ASA ethanol stock solutions were thawed once and then diluted 1 in 50 to working concentrations of 0.15 and 1.5 mg/mL in sterile saline, respectively. The combination NSAID treatment contained both 0.15 mg/mL INDO and 1.5 mg/mL ASA. Mice were treated by daily oral gavage with 20 µL/g body weight of respective working treatment solutions to achieve a final dose of 3 mg/kg/day INDO, 30 mg/kg/day ASA, or a combination of both 3 mg/kg/day INDO and 30 mg/kg/day ASA. These dosages were carefully selected to reflect clinically relevant human equivalent doses (∼170 mg and ∼17 mg per day, respectively) within their established therapeutic ranges. The matching vehicle (VEH) control solution consisted of an equivalent 4% ethanol in sterile saline. The first NSAID dose was administered to mice ∼5 min prior to BaCl_2_-induced skeletal muscle injury and then continued every 24 hours thereafter for the duration of the experiments. NSAIDs were administered daily shortly prior to onset of the dark phase.

### Muscle force testing

Isometric contractile properties of the TA muscle were measured *in situ* using the Aurora Scientific 1300A 3-in-1 Whole Animal System for Mice. Mice were anesthetized with 2% isoflurane, and the distal part of the TA muscle was isolated. The knee joint was pinned to the limb plate, and the distal TA tendon was tied to the lever arm of the force-test apparatus by a 4/0 silk suture (FST, 18020-40). The muscle was stimulated via platinum electrodes inserted through the overlying skin and fascia. Current and muscle length were adjusted to obtain supramaximal stimulation (0.2 ms pulse width) and optimal muscle length (*L*_o_), ensuring maximum isometric twitch force (*P*_t_). Sterile PBS was continuously applied to the exposed muscle to prevent drying. To determine the force-frequency relationship, the muscle was stimulated at 500 ms train durations at increasing frequencies (10–300 Hz) with 1-minute rest intervals between contractions. Maximum isometric tetanic force (*P*_o_) was defined as the highest force recorded during this protocol. Optimum fiber length (*L*_f_) was calculated by multiplying *L*_o_ by the TA-specific *L*_f_/*L*_o_ ratio of 0.6. The physiological CSA of the TA muscle was calculated as muscle mass divided by the product of *L*_f_ and the muscle density constant (1.06 mg/mm^3^). Specific tetanic force *sP*_o_ was calculated as *P*_o_/muscle CSA.

### Histology and immunofluorescence

Tissue cross-sections (10 µm) were cut from the mid-belly region of the TA muscle using a Leica CM1950 cryostat at –20°C. Muscle tissue sections were collected onto Super Frost Plus slides and air-dried at room temperature. Unfixed sections were used for hematoxylin and eosin (H & E) and muscle fiber type staining. Slides were fixed in 100% acetone for 10 minutes at −20°C before air-drying in preparation for staining of intramuscular immune cell populations. Prepared slides were blocked using either 10% normal goat serum (GS) (Invitrogen, 10000C) or with Mouse on Mouse (M.O.M.) IgG Blocking Reagent (Vector Laboratories, MKB22131) when mouse primary antibodies were used on mouse tissue samples. The slides were then incubated with primary antibodies prepared in either 10% GS in PBS or M.O.M protein diluent as appropriate overnight at 4°C. On the following day, the sections were washed 3 × 5 min in PBS and then incubated with appropriate Alexa Fluor-conjugated secondary antibodies (diluted 1:500 in PBS) for 1 h at room temperature. Slides were washed a further 3 × 5 min in PBS and then mounted with coverslips using MOWIOL Fluorescence Mounting Medium. Primary antibodies used include MyHC type I [Developmental Studies Hybridoma Bank (DSHB), BA-D5c, 1:100], MyHC type IIA (DSHB, SC-71c, 1:100), MyHC type IIB (DSHB, BF-F3c, 1:100), eMHC (DSHB, F1.652s, 1:20), Ly6G (GR1) (Bio-Rad, MCA2387, 1:50), CD68 (Bio-Rad, MCA1957, 1:200), CD206 (Bio-Rad, MCA2387, 1:50), and laminin (Abcam, ab7463, 1:200). Primary antibody staining was visualized with appropriate Alexa Fluor conjugated secondary antibodies (Invitrogen, 1:500 in PBS). DAPI (Invitrogen, Thermo Fisher Scientific, D21490, 2 μg/mL) was used to counterstain the cell nuclei. Stitched panoramic brightfield and fluorescent images of the entire TA muscle cross-section were captured using an automated fluorescent microscope (Echo Revolution) operating in upright configuration. Myofiber morphology, central nuclei fiber identification, and muscle fiber type profile was analyzed by high-throughput full automated image analysis using the MuscleJ 1.0.2 plugin for FIJI/ImageJ^75^.

### Muscle force testing

Isometric contractile properties of the TA muscle were measured *in situ* using the Aurora Scientific 1300A 3-in-1 Whole Animal System for Mice. Mice were anesthetized with 2% isoflurane, and the distal part of the TA muscle was isolated. The knee joint was pinned to the limb plate, and the distal TA tendon was tied to the lever arm of the force-test apparatus by a 4/0 silk suture (FST, 18020-40). The muscle was stimulated via platinum electrodes inserted through the overlying skin and fascia. Current and muscle length were adjusted to obtain supramaximal stimulation (0.2 ms pulse width) and optimal muscle length (*L*_o_), ensuring maximum isometric twitch force (*P*_t_). Sterile PBS was continuously applied to the exposed muscle to prevent drying. To determine the force-frequency relationship, the muscle was stimulated at 500 ms train durations at increasing frequencies (10–300 Hz) with 1-minute rest intervals between contractions. Maximum isometric tetanic force (*P*_o_) was defined as the highest force recorded during this protocol. Optimum fiber length (*L*_f_) was calculated by multiplying *L*_o_ by the TA-specific *L*_f_/*L*_o_ ratio of 0.6. The physiological CSA of the TA muscle was calculated as muscle mass divided by the product of *L*_f_ and the muscle density constant (1.06 mg/mm^3^). Specific tetanic force *sP*_o_ was calculated as *P*_o_/muscle CSA.

### Statistical analysis

Data are presented as the mean ± SEM, with raw data from each biological replicate displayed as dot plots on column graphs. Statistical analysis was performed in GraphPad Prism 10. For experiments with one independent variable with ≥3 levels, differences between experimental groups were tested by one-way ANOVA followed by pairwise Holm-Šídák post hoc tests. For experiments with two independent variables, between-group differences were tested by a two-way ANOVA followed by pairwise Holm-Šídák post hoc tests. P ≤ 0.05 was used to determine statistical significance.

## RESULTS

### Aspirin preserves *in vitro* myogenic potential through a COX dependent mechanism

We first evaluated the ability of treatment with various NSAIDs on the *in vitro* myogenic differentiation of murine C2C12 myoblasts. Treatment with common reversible and non-selective COX inhibitors including indomethacin (200 µM) and ibuprofen (1 mM) markedly suppressed myotube formation following 72 h of myogenic differentiation (**Fig. 1A**). This inhibitory effect was even most potent with COX-2 selective inhibitors, such as celecoxib (50 µM) and SC-236 (25 µM) (**Fig. 1A**). In sharp contrast, in our initial experiments ASA (2 mM) had no obvious discernible negative impact on myotube formation (**Fig. 1A**). Subsequent dose-response experiments highlighted a wide safety margin for ASA compared to non-ASA NSAIDs (**Fig. 1B**). Indomethacin had some statistically significant deleterious effects at doses as low as 25 µM (**Fig. 1C-E**). In contrast, ASA maintained myogenic capacity across a relatively broad range (**Fig. 1F**). Deleterious effects with ASA were only observed at overtly supraphysiological concentrations between 2 and 4 mM, most likely reflecting generalized cellular toxicity rather than targeted COX-mediated impairment of myogenesis (**Fig. 1H-J**). Crucially, the apparent preservation of myogenesis by lower doses of ASA appeared to be contingent upon its ability to interact with active COX enzymes (**Fig. 1K**). In co-treatment experiments, while ASA alone (2 mM) did not hinder differentiation, co-administration of this dose of ASA together with a moderate dose of indomethacin (200 µM) resulted in a markedly more severe blockade of myotube formation than either inhibitor alone (**Fig. 1L-N**). This suggests that the relatively protective profile of ASA treatment upon myogenesis *in vitro* is lost when the COX enzyme is competitively occupied by other reversible non-ASA NSAIDs such as INDO.

**Figure 1.**
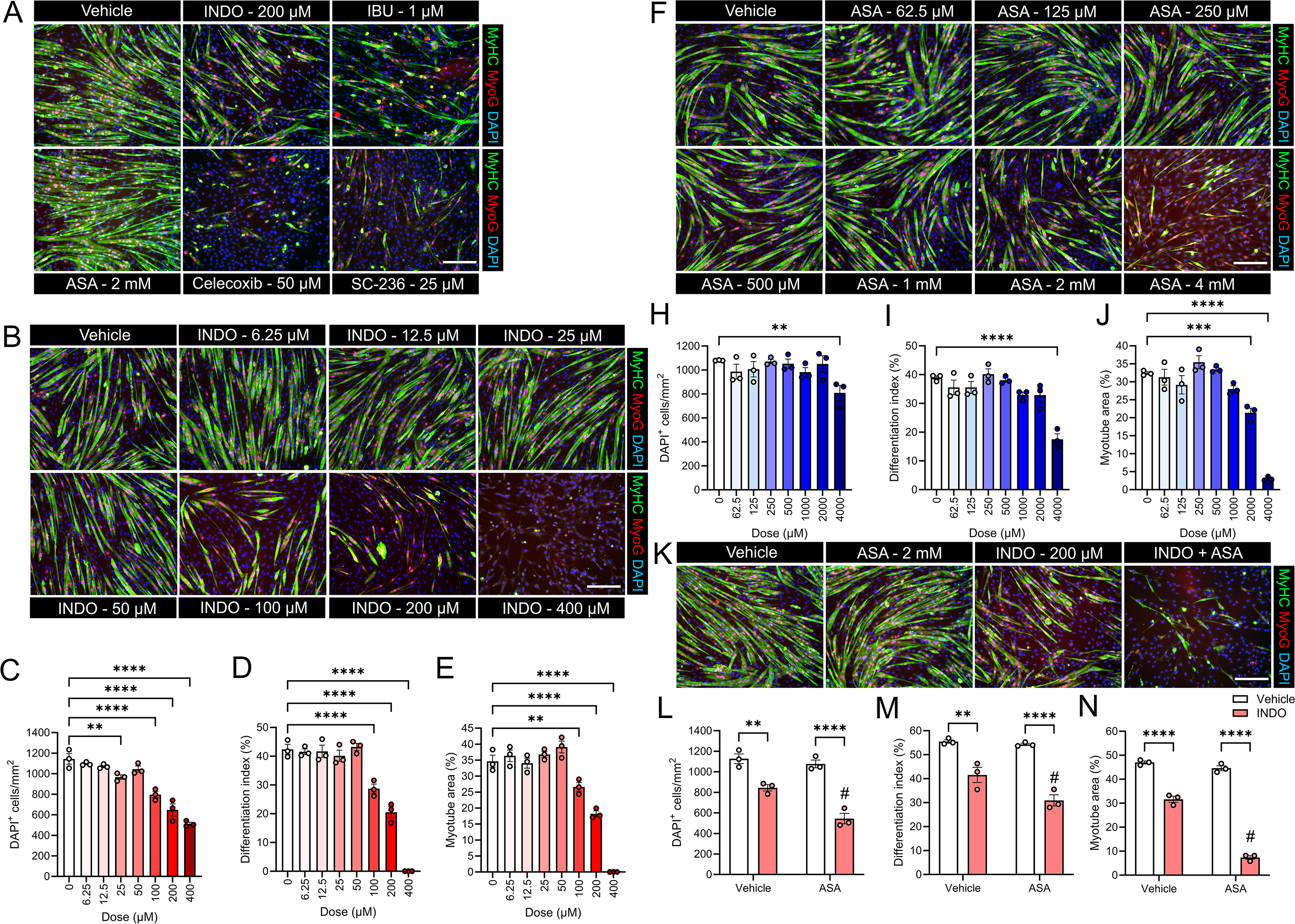
Aspirin and non-aspirin NSAIDs differentially impact *in vitro* myoblast differentiation and myotube formation. **A:** Representative immunofluorescence images of day 3 post-differentiation C2C12 myotubes that were treated during differentiation with various NSAIDs, including aspirin (ASA), indomethacin (INDO), ibuprofen (IBU), celecoxib, and SC-236 at the indicated concentrations. Cells were stained for myosin heavy chain (MyHC, green), myogenin (MyoG, red), and DAPI (nuclei, blue). **B:** Representative immunofluorescence images of day 3 post-differentiation C2C12 myotubes treated with doses of INDO ranging from 6.25 µM to 400 µM. **C-E**: Quantitative analysis of INDO-treated cells showing a dose-dependent decrease in DAPI^+^ cell density (**C**), differentiation index (%) (**D**), and myotube area (%) (**E**). **F:** Representative images of ASA-treated myotubes ranging from 62.5 µM to 4 mM. **H–J:** Quantification of DAPI^+^ cell density (**H**), differentiation index (%) (**I**), and myotube area (%) (**J**) in C2C12 cells receiving ASA treatment. **K:** Representative images comparing the effects of ASA (2 mM), INDO (200 µM), and the combination of INDO + ASA on C2C12 myotube morphology. **L-N:** Statistical comparison of cell density (**L**), differentiation index (%) (**M**), and myotube area (%) (**N**) across vehicle and treatment groups. Data are expressed as mean ± SEM. Statistical significance was determined by one-way ANOVA followed by Holm-Šídák post hoc tests. *, **, ***, and **** denotes p<0.05, p<0.01, p<0.001, and p<0.0001 between indicated groups, respectively. # denotes p<0.05 difference between ASA and vehicle groups.

### Aspirin promotes the resolution of acute skeletal muscle inflammation *in vivo* via a COX dependent mechanism

To assess the potential differential effects of ASA and INDO on acute skeletal muscle inflammation, we characterized the intramuscular immune landscape 24 hours post-injury (**Fig. 2A**). While total counts of infiltrating polymorphonuclear neutrophils (PMNs) (Ly6G^+^ cells) (**Fig. 2B**) and monocytes/macrophages (MΦs) (CD68^+^ cells) (**Fig. 2C**) were unchanged across treatment groups, the composition of the inflammatory infiltrate was fundamentally altered by ASA. Specifically, ASA significantly reduced the Ly6G^+^/CD68^+^ cell ratio, indicating an accelerated transition from PMN dominance toward monocyte/MΦ led resolution (**Fig. 2D**). Furthermore, ASA significantly increased the M1 (CD68^+^CD206^-^) to M2 (CD206^+^) MΦ ratio at this time point suggesting that ASA may promote the early recruitment of infiltrating inflammatory blood monocytes (CD68^+^CD206^-^ cells) that are required for the subsequent clearance of apoptotic PMNs and necrotic myofiber debris during the later stages of muscle regeneration (**Fig. 2E**). Crucially, these pro-resolving effects of ASA administration were abolished by INDO co-treatment, with the combination group remaining statistically indistinguishable from INDO alone (**Fig. 2E-F).** This suggests that INDO may act via competitive occupancy to block ASA from irreversibly acetylating the active site of COX-2. In theory, this could prevent the transition toward a pro-resolving phenotype, potentially by inhibiting the production of AT-SPMs.

**Figure 2.**
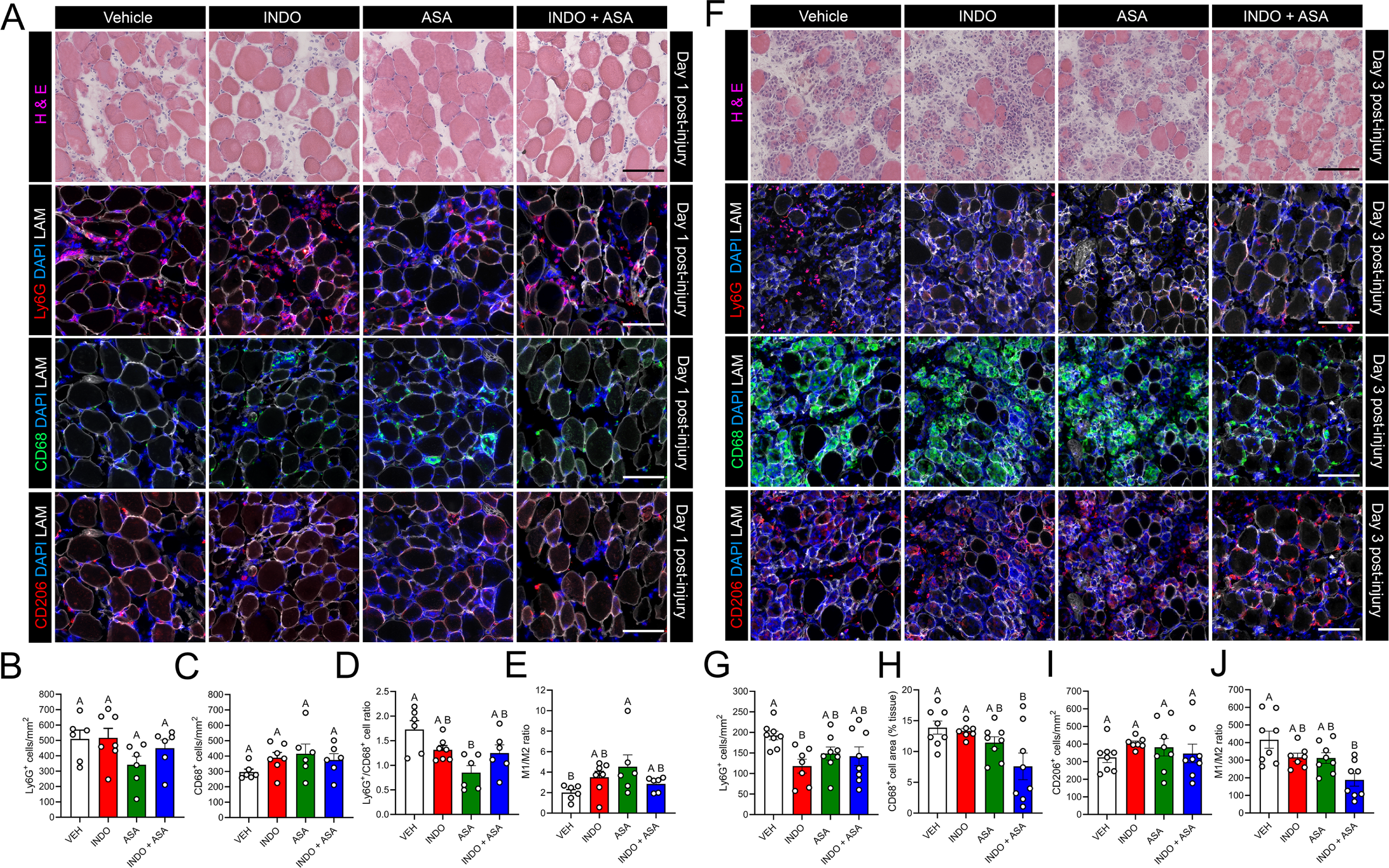
NSAID treatment modulates inflammatory cell infiltration following acute muscle injury. **A:** Representative histological (H & E) and immunofluorescence images of TA muscle cross-sections at day 1 post-injury following treatment with vehicle (VEH), indomethacin (INDO), aspirin (ASA), or a combination of INDO + ASA. Sections are stained for Ly6G (neutrophils, red), CD68 (total macrophages, green), CD206 (M2-like macrophages, red), with DAPI (nuclei, blue) and Laminin (LAM, white) to visualize fiber boundaries. **B-E**: Quantification of day 1 post-injury inflammatory markers including Ly6G^+^ cell density (**B**), CD68^+^ cell density (**C**), the ratio of Ly6G^+^ to total CD68^+^ cells (**D**), and the M1/M2 macrophage ratio (**E**). **F:** Representative H & E and immunofluorescence images at day 3 post-injury for the same treatment groups and markers. **G-J:** Quantitative analysis of inflammatory cell dynamics at day 3 post-injury, including Ly6G^+^ cell density (**G**), CD68^+^ cell area as a percentage of total tissue (**H**), CD206^+^ cell densities (**I**), and the M1/M2 ratio (**J**). All data are presented as mean ± SEM. Statistical significance was determined by one-way ANOVA followed by Holm-Šídák post hoc tests. Groups labeled with different letters are significantly different from one another, while groups sharing a common letter are not significantly different.

We further characterized skeletal muscle inflammation at 72 hours post-injury (**Fig. 2F**). At this point the INDO group showed significantly lower intramuscular PMNs (Ly6G^+^ cells) than vehicle (VEH) treated mice (**Fig. 2G**). Mice receiving ASA alone or a combination of both ASA and INDO showed an intermediate anti-inflammatory effect with intramuscular PMN counts not differing significantly from either the INDO or VEH groups (**Fig. 2G**). Large numbers of pro-inflammatory M1 MΦ (CD68^+^CD206^-^ cells) were seen infiltrating within the bodies of necrotic myofibers at this time-point. However, mice treated with a combination of INDO and ASA showed a marked inhibition of intramuscular infiltration of inflammatory M1 monocytes/MΦs (**Fig. 2H**). The absolute number of M2 MΦs (CD206^+^ cells) did not differ between experimental groups (**Fig. 2I**). Nevertheless, the ratio of M1 to M2 MΦ was significantly lower in mice receiving treatment with a combination of ASA + INDO when compared to VEH, indicating a blockade of recruitment of inflammatory blood monocytes and/or a persistent presence of resident M2-like MΦ within injured skeletal muscle (**Fig. 2J**).

### Indomethacin and aspirin have opposing effects on early indices of skeletal muscle regeneration

To assess the effect of ASA and INDO on early indices of myofiber regeneration, injured TA muscles were collected at 5 days following intramuscular BaCl_2_ injection. BaCl_2_ injury resulted in the appearance of numerous regenerating myofibers with characteristic centrally located myonuclei that expressed the developmental marker embryonic myosin heavy chain (eMHC) (**Fig. 3A**). The absolute number of eMHC^+^ myofibers did not differ across groups, indicating no effect INDO, ASA, or a combination of INDO + ASA on the initial formation of new myofibers in injured muscle (**Fig. 3B**). Nevertheless, mice receiving treatment with INDO alone did display a significant reduction in mean eMHC^+^ myofiber cross-sectional area (CSA) when compared to the VEH group (**Fig. 3C**). In contrast, the mean CSA of the eMHC^+^ myofiber population in ASA-treated mice did not differ from VEH and showed a statistical trend toward being larger than those in the INDO group. Surprisingly, mice receiving the ASA + INDO combination also exhibited larger eMHC^+^ fibers than INDO and were further statistically indistinguishable from the ASA-only and VEH groups (**Fig. 3C**). Further analysis of the percentage frequency distribution of regenerating myofiber size revealed that INDO treatment shifted the regenerating population toward a presence of smaller myofibers, with a greater proportion of fibers in the 50-300 µm^2^ range and a reduced proportion in the 450-650 µm^2^ range (**Fig. 3D**). In contrast, the percentage frequency distribution of eMHC^+^ fibers did not differ between VEH and ASA treated mice at any CSA bins (**Fig. 3E**). The frequency distribution of regenerating myofiber CSA was generally similar between mice treated with VEH and a combination of both ASA + INDO, except for a greater proportion of fibers in the 50-100 µm^2^ range and a lower proportion of fibers in the 400-450 µm^2^ range in mice receiving ASA + INDO (**Fig. 3F**). These results show that ASA and INDO have differential effects on myofiber regeneration *in vivo*. Specifically, unlike INDO, ASA appears to uniquely preserve the growth of newly formed myofibers. Furthermore, surprisingly ASA also appears to partially protect against the growth-inhibitory effects of INDO.

**Figure 3.**
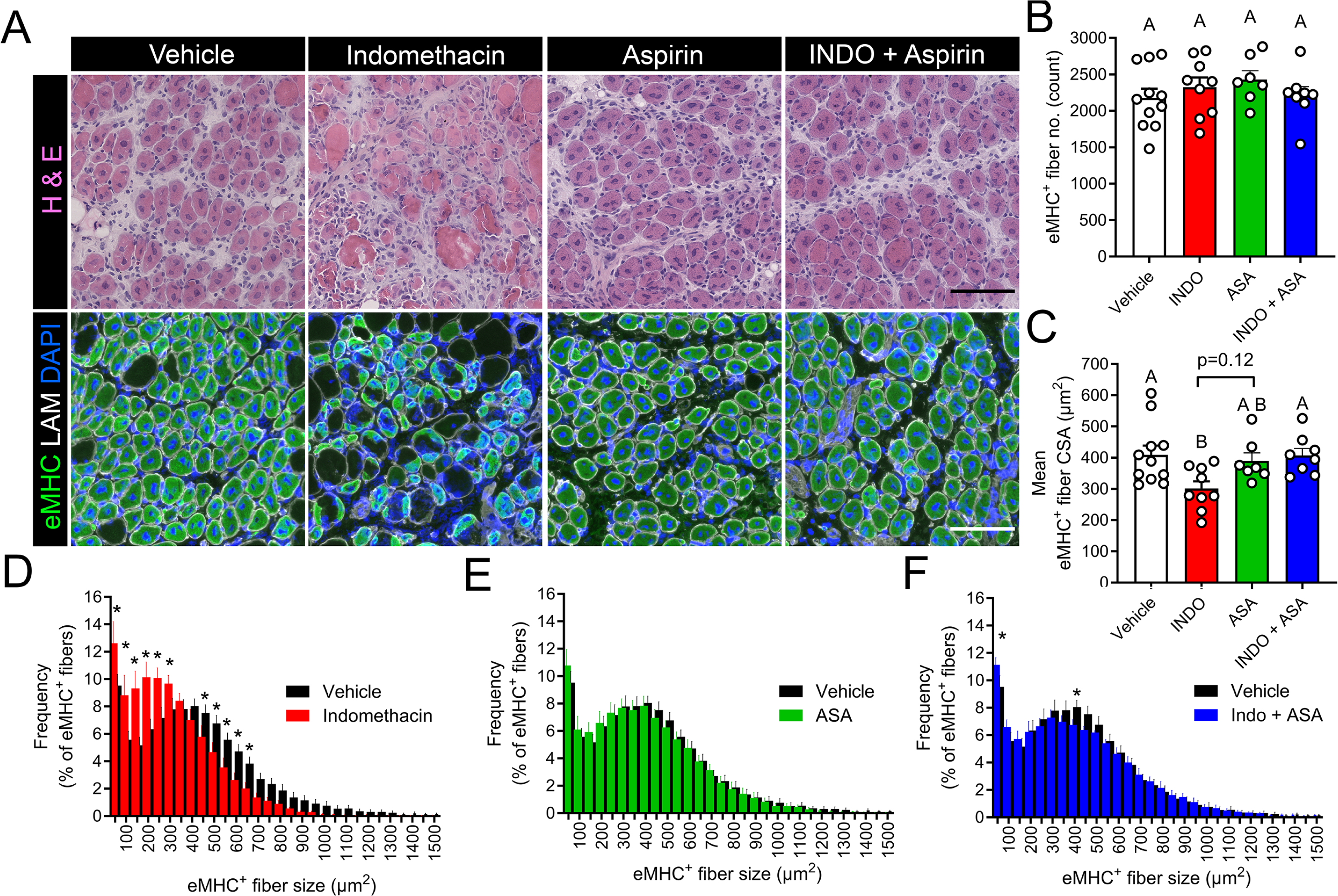
Treatment with indomethacin, but not aspirin, reduces the size of regenerating myofibers following acute muscle injury. **A:** Representative histological (H & E, top) and immunofluorescence (bottom) images of regenerating muscle cross-sections at day 5 post-injury. Mice were treated via daily oral gavage with vehicle (VEH), indomethacin (INDO), aspirin (ASA), or a combination of both INDO + ASA. Immunofluorescence highlights embryonic myosin heavy chain (eMHC, green) to identify regenerating fibers, with Laminin (LAM, white) delineating fiber boundaries and DAPI (blue) labeling nuclei. **B:** Total count of eMyHC+ regenerating fibers across treatment groups. **C:** Mean cross-sectional area (CSA) of the regenerating eMHC^+^ myofiber population. **D-F:** Frequency distribution plots showing the percentage of eMHC^+^ fibers across different size ranges for mice treated with INDO vs. VEH (**D**), ASA vs. VEH (**E**), and Indo + ASA vs. VEH (**F**). Data are presented as mean ± SEM. **A-C:** Statistical significance was determined by one-way ANOVA followed by Holm-Šídák post hoc tests. Groups labeled with different letters are significantly different from one another, while groups sharing a common letter are not significantly different. **D-F:** Statistical significance was determined by unpaired two-tailed t-tests. * denotes p<0.05 difference between VEH and NSAID treated mice.

### Aspirin promotes functional recovery of absolute muscle contractile capacity while indomethacin exacerbates strength deficits

To assess functional recovery of muscle strength, we analyzed *in situ* isometric force-frequency relationships of the TA muscle at day 14 post-injury. Two-way ANOVA revealed a significant treatment × frequency interaction effect for absolute muscle force generating capacity (mN). While muscle strength remained significantly depressed in the VEH, INDO, and INDO + ASA groups when compared with uninjured control (CON) mice at frequencies of 80-300 Hz, ASA-treated mice achieved force levels across all frequencies (10-300 Hz) that were statistically indistinguishable from CON (**Fig. 4A**). Notably, INDO treatment further exacerbated absolute muscle strength deficits when compared to injured mice receiving either VEH or ASA treatment. Integration of the area under the curve (AUC) confirmed that while all injured groups exhibited a net deficit in total contractile capacity compared with uninjured CON (**Fig. 4B**), that ASA-treated mice displayed superior overall strength relative to the VEH group. Conversely, INDO treatment resulted in the lowest overall force production with an AUC significantly lower than mice receiving either VEH or ASA. Crucially, any potential therapeutic benefit of ASA was completely abolished in the INDO + ASA group, which statistically mirrored the INDO-only group (**Fig. 4B**). These data suggest that the modest, yet statistically significant functional benefits of ASA on recovery of absolute force production requires COX-2 activity which is blocked by the competitive binding of INDO throughout the 14-day recovery.

**Figure 4.**
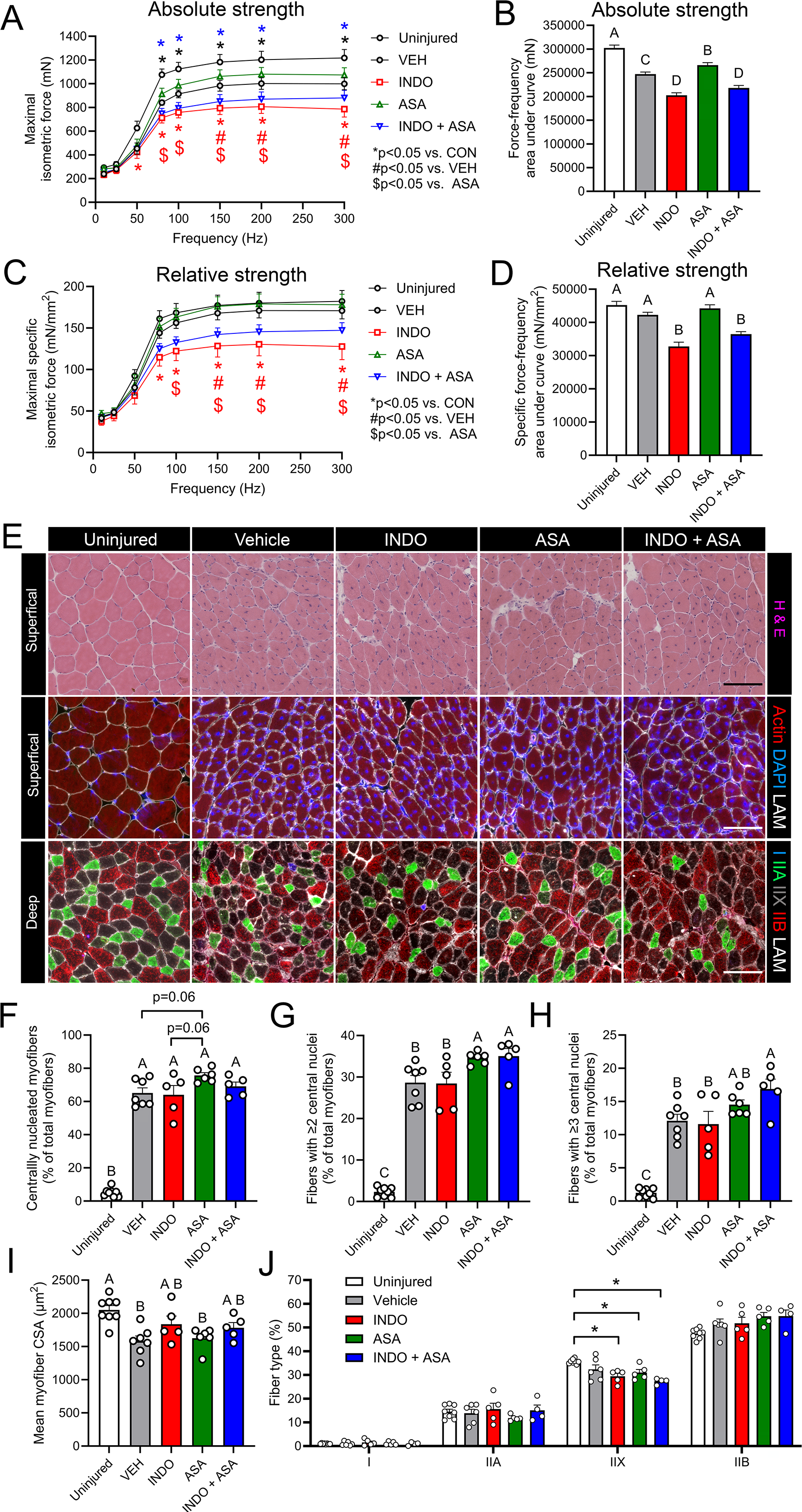
Non-aspirin NSAIDs impair functional recovery and myofiber maturation following acute muscle injury, while aspirin promote recovery of absolute muscle strength. **A:** Force-frequency curves showing maximal isometric tetanic force generated by the TA muscle *in situ* at day 14 post-injury. **B:** Total area under the force-frequency curve (absolute muscle strength). **C-D:** Specific force-frequency curves (**C**) and specific force-frequency area under the curve (**D**) (relative muscle strength) normalized to muscle size. **E:** Representative histological and immunofluorescence images of muscle cross-sections. Top: H & E staining; Middle: Actin (red), DAPI (blue), and Laminin (LAM, white); Bottom: Fiber type staining for type I (blue), type IIA (green), type IIX (black), and type IIX (red), and laminin (white). **F-H:** Quantification of regenerative maturation markers including the percentage of centrally nucleated myofibers (**F**), fibers with ≥2 central nuclei (**G**), and fibers with ≥3 central nuclei (**H**). **I-J:** Mean myofiber cross-sectional area (CSA) (**I**) and muscle fiber type profile (I, IIA, IIX, IIB) (**J**). Data are presented as mean ± SEM. For force-frequency curves statistical significance was determined by two-way ANOVA followed by Holm-Šídák post hoc tests. * denotes p<0.05 vs. uninjured control mice (CON), # denotes p<0.05 vs. injured mice receiving VEH, and $ denotes p<0.05 vs. injured mice receiving ASA. For bar graphs, different letters indicate significant differences (p<0.05) by one-way ANOVA with Holm-Šídák post hoc tests.

### Complete COX inhibition by indomethacin, but not COX acetylation by aspirin, persistently impairs myofiber contractile quality

To determine if the effects of ASA and INDO on strength recovery was driven by changes in muscle size or intrinsic fiber quality, we calculated specific muscle force (mN/mm²) by normalizing absolute tension to the physiological muscle CSA. Two-way ANOVA revealed a significant treatment × frequency interaction effect. Injured mice treated with INDO displayed significantly lower specific muscle force when compared with both uninjured CON mice or injured VEH and ASA groups (**Fig. 4C**). This functional impairment was particularly evident at higher stimulation frequencies (100–300 Hz). Mice co-treated with INDO + ASA were statistically indistinguishable from those receiving INDO alone. Integration of the force-frequency relationship (AUC) confirmed that VEH and ASA groups achieved specific force levels indistinguishable from uninjured CON (**Fig. 4D**). In contrast, muscles from mice treated with either INDO alone or a combination of both INDO + ASA remained functionally depressed. These data demonstrate that while ASA preserves the intrinsic contractile quality of regenerating myofibers, potent non-selective COX inhibition by INDO results in persistent functional deficits in specific muscle force.

### ASA-induced myonuclear accretion is independent of COX-mediated functional recovery

To investigate the cellular basis underlying the differences in skeletal muscle contractile force, we quantified the percentage of regenerating centrally nucleated fibers (CNFs) and their relative density of their centrally located myonuclei in muscle samples collected on day 14 post-injury. While the total proportion of CNF was markedly elevated in all injured groups relative to uninjured CON, ASA-treated mice tended to have a greater proportion of regenerating myofibers when compared to the VEH or INDO groups (p=0.06) (**Fig. 4F**). ASA-treated mice also exhibited higher overall myonuclear accretion as indicated by a significantly greater proportion of fibers containing ≥2 centrally located nuclei (**Fig. 4G**), and a similar upward trend observed for fibers with ≥3 central nuclei (**Fig. 4H**). Intriguingly, unlike the recovery of muscle strength, this apparent increase in myonuclear accretion in ASA-treated mice was clearly not abolished in the INDO + ASA group. Rather mice treated with a combination of INDO + ASA displayed a greater proportion myofibers with both ≥2 central nuclei (**Fig. 4G)** and ≥3 central nuclei (**Fig. 4H)** when compared to the VEH group. Despite the significant growth delays observed in the INDO group at day 5 post-injury (see **Fig. 3**), mean myofiber CSA at day 14, was surprisingly uniform across all injured groups (**Fig. 4I**). These data suggest that ASA may promote satellite cell fusion and myonuclear accretion through a distinct COX-independent mechanism that persists even in presence of INDO co-treatment. However, the failure of the INDO + ASA group to achieve functional recovery of muscle strength following 14-days after injury, despite this enhanced multinucleation, underscores that COX-dependent resolution signals appear to be essential for translating increased myoblast fusion into intrinsic improvements in myofiber contractile quality. Further analysis of muscle fiber type profile revealed that mice treated with INDO, ASA, or a combination of INDO + ASA displayed a significantly reduced proportion of type IIX muscle fibers relative to uninjured CON mice (**Fig. 4J**). Nevertheless, the fiber type profile of regenerating muscle was overall similar between treatment groups and thus unlikely to explain the differences seen in contractile muscle force.

## DISCUSSION

In the current study we assessed the potential unique impact of ASA on inflammation and regeneration following acute skeletal muscle injury. Consistent with prior studies suggesting that non-ASA NSAIDs can interfere with muscle repair^4^, INDO blunted *in vitro* myogenesis, reduced early regenerating myofiber size *in vivo*, and impaired recovery of muscle strength in mice following BaCl_2_-induced injury. In contrast, ASA was relatively permissive for *in vitro* myogenesis, accelerated the resolution of acute muscle inflammation, did not interfere with myofiber regeneration, and improved recovery of muscle strength. The ability of ASA to both hasten the resolution of muscle inflammation and promote functional recovery were abolished by co-treatment with INDO. Surprisingly, however, the apparent beneficial effects of ASA on myonuclear accretion were not affected by INDO co-treatment. Overall, these data show that unlike other non-ASA NSAIDs which can have deleterious effects on skeletal muscle regeneration, ASA is relatively safe in terms of its impact on muscle regenerative capacity and rather hastens the natural resolution of acute inflammation leading to improved functional recovery of muscle strength via a COX-dependent pro-resolving mechanism We found that a range of common competitive and reversible non-selective COX inhibitors including INDO and IBU had marked deleterious effects on the ability of C2C12 myoblasts to undergo myogenic differentiation and fusion to form mature myotubes *in vitro*. This finding is consistent with numerous prior studies by us and others that have reported marked negative effects of non-selective NSAIDs such as INDO, IBU, and naproxen sodium on *in vitro* myotube formation^48,51,66,76,77^. Interestingly, one recent study found that while consistent with our data that treatment with INDO at a dose of 200 µM reduced the fusion index of human primary muscle cells, that similar doses of IBU did not have such negative effects^41^. However, this result might potentially be explained by the dose of IBU tested as we generally find that relatively higher doses of ibuprofen (e.g., ≥500 µM are required to observe obvious deleterious effects on C2C12 myotube formation when compared to other non-ASA NSAIDs^66^.

Consistent with prior reports, we further found that the COX-2 selective inhibitors celecoxib and SC-236 were even more potent inhibitors of myogenesis than non-selective NSAIDs^48,49,51,66^. The marked negative effects of celecoxib on myotube formation are consistent with a recent study in human primary myoblasts^57^. Interestingly, it was concluded by the authors in this recent study that the deleterious effects of celecoxib on myogenesis were independent of COX-2, in part, because similar negative effects were not observed with a distinct COX-2 inhibitor NS-398^57^. Nevertheless, several prior reports by us and other groups have indeed reported marked negative effects of NS-398 *in vitro* muscle cell growth and development^48,49,51,66^. We generally do find that relatively higher doses of NS-398 (≥50 µM) are required to observe obvious deleterious effects on myotube formation when compared to other COX-2 selective NSAIDs^48,66^. In support of the argument that the negative effects of COX-2 selective inhibitors including celecoxib, SC-236, and NS-398 on *in vitro* myogenesis may indeed be directly attributable to reduced COX-2 activity, siRNA mediated knockdown of COX-2 in C2C12 myoblasts also interferes with myogenic differentiation and fusion of C2C12 myoblasts^78^. Furthermore, primary myoblasts isolated from COX-2 knockout (COX-2^-/-^) mice display a markedly impaired ability to fuse to form mature myotubes^79^. Most strikingly, such impaired fusion of COX-2^-/-^ myoblasts can be reversed by repletion of major COX-derived eicosanoids including PGE_2_ or PGF_2α_^79^. Overall, this body of literature provides convincing evidence that COX-2 mediated eicosanoid biosynthesis is an indispensable event for successful myogenesis *in vitro*.

Surprisingly, despite ASA being one of the most used non-selective NSAIDs worldwide to our knowledge no prior study has tested the effect of ASA specifically on myogenesis *in vitro*. In our initial studies we did not observe any obvious deleterious effects of treatment of differentiating C2C12 myoblasts with ASA at doses as high as 2 mM. Nevertheless, in follow-up dose-response experiments we did observe some reproducible negative effects of aspirin on myotube formation when administered in the 2-4 mM range. The relatively lower inhibitory dose of ASA observed in these subsequent dose-response experiments may be due to the added cellular stress of the substantial concentration of ethanol vehicle (0.8%) required to test ASA doses as high as 4 mM given the solubility of ASA in this solvent (∼80 mg/mL or ∼450 mM). When tested in parallel under these same conditions, INDO showed some negative effects at doses as low as ∼25 µM and displayed virtually complete blocked myotube formation at doses of ∼400 µM. Interestingly, doses of ASA which had no obvious negative effects on myogenesis when administered alone (∼1-2 mM) did clearly exacerbate the negative effect of moderate doses of INDO (∼200 µM). This suggests that ASA-induced acetylation of COX-1 and 2 can indeed have negative effects on myogenesis when the overall enzymatic activity of COX is simultaneously blocked by other non-selective NSAIDs, presumably via further lowered prostaglandin biosynthesis. An alternative explanation may be that when INDO competitively binds to and prevents ASA from interacting with the COX enzymes, ASA has other COX-independent negative effects on myogenesis like those observed in response to extreme doses of aspirin alone (e.g., 4 mM). Overall, these findings show that ASA appears to be relatively safe compared to other non-aspirin NSAIDs in terms of its ability to not interfere with myogenesis *in vitro* when administered at physiologically relevant doses. Nevertheless, the permissive effects of lower doses of ASA upon myogenesis *in vitro* do appear to be dependent on intact COX activity.

We found that oral gavage with a single preemptive oral dose of INDO (3 mg/kg) at the time of BaCl_2_-induced muscle injury did not influence absolute intramuscular numbers of PMNs, MΦs, or the ratio of PMNs/MΦ at 1-day post-injury. This finding contrasts with several prior studies that have reported that treatment with either non-ASA NSAIDs^80–82^ or COX-2 specific inhibitors^49,53,56^ could reduce overall leukocyte infiltration of injured skeletal muscle. Nevertheless, interestingly our INDO group data is consistent with several other prior studies that have reported that NSAID treatment perhaps suprisingly failed to obviously impact overall muscle leukocyte infiltration^39,43,68,83–85^. Unlike INDO, we found that a single preemptive oral dose of ASA (30 mg/kg) did tend to decrease the absolute intramuscular numbers of PMNs, while also tending to increase absolute presence of MΦ, ultimately resulting in a significantly decreased PMN/MΦ ratio at day 1 post-injury. Additionally, we found that the ratio of total monocytes/MΦ (CD68^+^ cells) to M2-like MΦ (CD206^+^ cells) was increased at day 1 post-injury in mice receiving ASA treatment. Collectively, this suggests that ASA may promote recruitment of blood monocytes to the site of injury while simultaneously limiting PMN infiltration and/or stimulating PMN clearance. As mentioned above, this effect of ASA appears to be generally opposite prior studies of non-ASA NSAIDs which have generally been found to either reduce^80–82^, or fail to influence^39,43,68,83–85^, blood monocyte recruitment to injured muscle. Both the reduced PMN/MΦ and increased total MΦ/M2 MΦ ratio at day 1 post-injury in the ASA group were not observed in mice receiving combination treatment of ASA + INDO. Thus, while we did not find strong evidence to suggest that INDO alone delayed resolution of muscle inflammation following BaCl_2_-induced injury, it nevertheless did appear to interfere with the ability of ASA to promote timely inflammation-resolution at day 1 post-injury.

Mice treated with INDO showed fewer intramuscular PMNs with a similar trend observed for ASA at day 3 post-injury. This finding suggests that both INDO and ASA can reduce PMN presence at this time-point, but that perhaps surprisingly the combined impact of INDO + ASA on intramuscular PMNs does not appear to be cumulative. This may suggest that INDO and ASA act to reduce PMN numbers by different mechanisms. Neither INDO or ASA alone reduced total intramuscular MΦ numbers at this time-point. Nevertheless, treatment with a combination of both ASA + INDO did markedly reduce total MΦ infiltration of injured muscle. Overall, this finding is consistent with our *in vitro* data showing that while physiological doses of ASA alone do not impair myogenesis, that in the presence of non-selective COX inhibition by INDO that the impact of ASA with regard to inflammatory monocyte/MΦ infiltration is also cumulative. Overall, this finding is consistent with some prior published studies suggesting that NSAIDs may reduce blood monocyte/M1 MΦ recruitment to injured muscle^80–82^. On the other hand, the lack of obvious effect of INDO or ASA alone on intramuscular numbers of MΦ is indeed consistent with many numerous prior studies that have failed to observe a show suppressive effect of non-ASA NSAIDs on monocyte/MΦ infiltration following skeletal muscle injury^39,43,68,83–85^.

We found that daily oral gavage with INDO (3 mg/kg/day) did not influence the total number of regenerating (eMHC^+^) myofibers at day 5 post-injury but nevertheless did reduce regenerating fiber CSA at this time-point. This finding is consistent with some prior studies that have reported a negative effect of the non-selective NSAIDs such as piroxicam^80^ and IBU^39^, as well as COX-2 specific inhibitors such as NS-398^49^ and SC-236^53^ on regenerating muscle fiber size following acute muscle injury. Consistently, non-aspirin NSAIDs and COX-2 specific inhibitors have both been shown to blunt normal increases in myofiber CSA in models of modified muscle use including exercise^38^, overload-induced hypertrophy^37,50^, and muscle regrowth following disuse atrophy^44,54^. Like INDO, daily gavage with ASA did not affect the number of regenerating myofibers. But, unlike INDO, we found no evidence that ASA had any negative effect on eMHC^+^ myofiber CSA at day 5 post-injury. Furthermore, mice receiving combinational treatment with INDO + ASA also showed no defects in regenerating muscle fiber number or size. Collectively, these data show that consistent with our *in vitro* data, ASA is relatively safe regarding the growth and development of regenerating muscle fibers during the early stages of recovery following muscle injury. Surprisingly, however, ASA treatment appears to rescue the deleterious effects of *in vivo* INDO treatment on early regenerating muscle fiber size. This suggests that ASA may act through some other COX-independent mechanism to stimulate myofiber growth even when COX activity is blocked by INDO. Notably, this contrasts with our *in vitro* data showing a clear cumulative negative effect of ASA on myotube formation in cells receiving INDO co-treatment. These contrasting findings could be related to differential responses between C2C12 cells grown under normal conditions and animal models in early phase of recovery following muscle injury.

We further observed that daily oral gavage with the non-selective NSAID INDO reduced recovery of absolute and specific contractile force generating capacity of the TA muscle *in situ* measured at day-14 post-injury. The apparent negative effect of INDO on functional recovery of muscle strength in the current study is consistent with some prior reports of impaired functional recovery in animals treated with non-selective non-aspirin NSAIDs including flurobiprofen^7^ or INDO^86^. Nevertheless, several other prior studies have failed to observe an obvious negative effect of non-aspirin NSAIDs including INDO^87^ and piroxicam^88^, the dual COX/5-LOX inhibitor licofelone^84^, or the COX-2 specific inhibitor mavacoxib^89^ on recovery of muscle strength in animal models of muscle injury. We found that unlike INDO, ASA treatment modestly increased absolute strength and neither enhanced nor perturbed recovery of specific muscle force. Mice receiving combinational treatment with both INDO + ASA also showed impaired functional recovery like those receiving INDO alone. Therefore, the modest yet statistically significant benefit of ASA on recovery of absolute muscle strength appears to require functional COX activity. Furthermore, ASA appears to have no functional benefits in the presence of non-selective COX inhibition despite the apparent ability of ASA to rescue early impairments in regenerating myofiber CSA induced by INDO.

In contrast to our observations of a deleterious effect of INDO treatment on eMHC^+^ myofiber CSA at day 5 post-injury we did not observe a persistent reduction in centrally nucleated muscle fiber CSA at day-14 of recovery from muscle injury. In fact, mice receiving INDO treatment tended to have slightly larger myofiber CSA than those receiving VEH treatment. Despite this, mice treated with INDO were clearly functionally impaired at this time-point based on *in situ* measurements of TA muscle strength. Consistent with this finding, a prior study also found that the non-aspirin NSAID flurbiprofen actually increased mean myofiber CSA at 28-days following eccentric contraction-induced skeletal muscle injury in rabbits despite an obvious deleterious effect of this NSAID on muscle strength at this same time-point^86^. Interestingly, while myofiber CSA was increased in rabbits receiving flurbiprofen treatment in this study, myofibril diameter was simultaneously reduced in the NSAID group^86^. Consistently, in the current study we found that while INDO not only reduced absolute strength (mN) at day-14 post-injury, but also impaired recovery of specific muscle force (mN/mm^2^) indicative of potential negative effect on intrinsic myofiber quality. Collectively, these data suggest that non-ASA NSAIDs may impair recovery of muscle function even when obvious defects at the histological level are not observed. This finding may potentially explain some of the conflicting reports in the literature that have failed to observe obvious deleterious effect of non-ASA NSAIDs on histological indices of skeletal muscle regeneration^43,87,88,90^.

We did not observe any positive or negative effects of ASA on regenerating muscle fiber CSA at day-14 post-injury. Nevertheless, interestingly ASA treated mice did tend to show an increased proportion of centrally nucleated myofibers at day 14 post-injury that was not observed in those receiving INDO. Moreover, ASA treatment enhanced cell fusion and myonuclear accretion of regenerating fibers as indicated by an increased proportion of muscle fibers with containing two or more centrally located myonuclei. Interestingly, these apparent beneficial effects of ASA were observed in mice receiving combination treatment with both ASA + INDO. This finding suggests that ASA uniquely promotes maturation of regenerating myofibers which may explain the benefits of ASA on recovery of absolute muscle strength. Surprisingly however, while this apparently enhanced fiber maturation was also observed in mice receiving combination treatment with ASA and INDO, it did not appear to translate to a functional benefit in terms of recovery of muscle strength.

Several limitations of this study warrant consideration. First, we did not directly measure intramuscular concentrations of bioactive AT-SPMs (e.g., AT-LXs, AT-RvDs, AT-NPD1, or RvEs) or their respective biosynthetic intermediates [15(R)-HETE, 17(R)-HDoHE, and 18(R)-HEPE]. Consequently, while our findings align with the known biochemical ‘switch’ of aspirin-acetylated COX-2, the specific molecular drivers of the observed pro-regenerative effects remain to be definitively identified. However, the observation that co-treatment with INDO, a potent inhibitor of both COX-1 and −2, reversed the beneficial effects of ASA on inflammatory resolution and muscle strength provides strong pharmacological evidence that these outcomes are indeed dependent on a COX-mediated pathway. Second, our *in vivo* experiments utilized a single dose of 30 mg/kg/day ASA. While this dose is optimized in rodents to trigger AT-SPM production without overt toxicity, representing a human equivalent dose of approximately 170 mg daily, we cannot exclude the possibility that higher, anti-inflammatory doses of ASA might yield different outcomes on myofiber recovery. Furthermore, while BaCl₂-induced injury provides a highly reproducible model of muscle regeneration, the pro-resolving impact of ASA may vary in clinical scenarios involving different injury kinetics, such as muscle strains or crush injuries. Finally, this study was conducted exclusively in female mice; given the known influence of sex hormones on both lipid metabolism, inflammatory resolution, and skeletal muscle regenerative ability, further research is required to determine if these beneficial effects are sex-dependent.

## CONCLUSIONS

In conclusion, our study demonstrates that ASA is unique among the NSAID class, as it does not impair, and may even facilitate skeletal muscle regeneration and functional recovery. While traditional non-selective and COX-2 selective NSAIDs like INDO markedly inhibit myogenesis, reduce early myofiber growth, and impair long-term functional recovery, physiological doses of ASA avoid these regenerative deficits. Mechanistically, we identified a novel divergence in ASA’s effects: while its benefits to the inflammatory environment and contractile strength are COX-dependent and were neutralized by co-treatment with INDO, the ability of ASA to promote myonuclear accretion and myofiber maturation surprisingly persisted even under competitive COX inhibition. These findings suggest that ASA utilizes both COX-dependent pro-resolving pathways and unique COX-independent mechanisms to support muscle repair. Collectively, our results challenge the long-standing clinical assumption that all NSAIDs uniformly impair muscle repair and suggest that ASA may represent a superior therapeutic strategy for managing musculoskeletal injuries without compromising the intrinsic quality or functional capacity of the regenerating tissue.

## ACKNOWLEDGEMENTS

The authors thank Shihuan Kuang (Duke University, formerly of Purdue University) for laboratory support and helpful discussions. The MF20 (developed by Fischman, D.A.) and F5D (developed by Wright, W.E.) monoclonal antibodies were obtained from the Developmental Studies Hybridoma Bank (DSHB), created by the NICHD of the NIH and maintained at The University of Iowa, Department of Biology, Iowa City, IA 52242.

## AUTHOR CONTRIBUTIONS

**Xinyue Lu:** Methodology, Formal analysis, Investigation, Writing – review & editing, Visualization. **Hamood Rehman:** Methodology, Software, Formal analysis, Investigation, Writing – review & editing, Visualization. **Amelie Sercu:** Investigation, Formal analysis, Writing – review & editing. **James F. Markworth:** Conceptualization, Methodology, Software, Formal analysis, Investigation, Resources, Writing – original draft, Writing – review & editing, Visualization, Supervision, Project administration, Funding acquisition. All authors approved the final draft and take responsibility for the data’s accuracy.

